# Corticostriatal Projections of Macaque Area 44

**DOI:** 10.1101/2020.10.23.352617

**Authors:** Cole Korponay, Eun Young Choi, Suzanne N. Haber

**Affiliations:** Basic Neuroscience Division, McLean Hospital, Belmont, MA; Department of Psychiatry, Harvard Medical School, Cambridge, MA; Department of Neurosurgery, Stanford University, Stanford, CA; Department of Pharmacology and Physiology, University of Rochester School of Medicine, Rochester, NY

**Keywords:** circuitry, frontostriatal, neuroanatomy, tract-tracing, orofacial motor control, area 44, area 6VR

## Abstract

Ventrolateral frontal area 44 is implicated in inhibitory motor functions and facilitating prefrontal control over vocalization. Yet, the corticostriatal circuitry that may contribute to area 44 functions is not clear, as prior investigation of area 44 corticostriatal projections is limited. Here, we used anterograde and retrograde tracing in macaques to map the innervation zone of area 44 corticostriatal projections, quantify their strengths, and evaluate their convergence with corticostriatal projections from non-motor and motor-related frontal regions. First, terminal fields from a rostral area 44 injection site were found primarily in the central caudate nucleus, whereas those from a caudal area 44 injection site were found primarily in the ventrolateral putamen. Second, amongst sampled striatal retrograde injection sites, area 44 input as a percentage of total frontal cortical input was highest in the ventral putamen at the level of the anterior commissure. Third, area 44 projections converged with both orofacial premotor area 6VR and other motor related projections (in the putamen), and with non-motor prefrontal projections (in the caudate nucleus). These findings support the role of area 44 as an interface between motor and non-motor functional domains, possibly facilitated by rostral and caudal area 44 subregions with distinct corticostriatal connectivity profiles.

## INTRODUCTION

The striatum receives dense projections from all regions of the frontal cortex[1–3]. Parsing this complex projection system is key to understanding how different cortical regions influence the output of the striatum. As such, a major focus of neuroanatomical investigations has been to determine where in the striatum different frontal cortical regions project to[1], the relative strengths of these projections[4], and how the convergence of projections from different regions [5] integrates diverse functional domains [6]. Invasive tract-tracing studies in non-human primates have proven invaluable for addressing these questions. Here, we use these methods to characterize the frontostriatal projections of macaque ventrolateral frontal area 44.

In the human brain, area 44 is implicated in speech production and in inhibitory motor control[7–9]. Similarly, area 44 in the macaque is involved in the control of vocalizations[10, 11], and there is some evidence for macaque area 44 involvement in broader inhibitory motor control [12]. Compared to other frontal cortical regions, the macaque homologue of human area 44 has been demarcated only relatively recently[11, 13, 14] (**Fig. 1a**). Comparative cytoarchitectonic analyses identified a narrow dysgranular region in the fundus of the inferior arcuate sulcus that is cytoarchitectonically distinct from the adjacent agranular premotor area 6VR (also referred to as F5[15]), and which resembles the cytoarchitectonic properties of area 44 in the human[11, 13]. Furthermore, electrophysiological data indicate that macaque area 44 is functionally distinct from the adjacent premotor cortex. In a study of macaque vocalization, area 44/45 neurons were found to discharge before vocal onset, whereas ventral premotor neurons discharged mostly concurrently with vocal onset[16]. Collectively, these data have prompted the conceptualization of area 44 as a transition region between the adjacent ventrolateral prefrontal and orofacial premotor cortices[10]. Retrograde labeling data, which demonstrates reciprocal connections between area 44 and both adjacent vlPFC area 45 and adjacent premotor area 6VR, further supports this notion[17, 18]. Functionally, this may position area 44 to mediate higher-order prefrontal control over orofacial motor activity[10].

**Figure 1:**
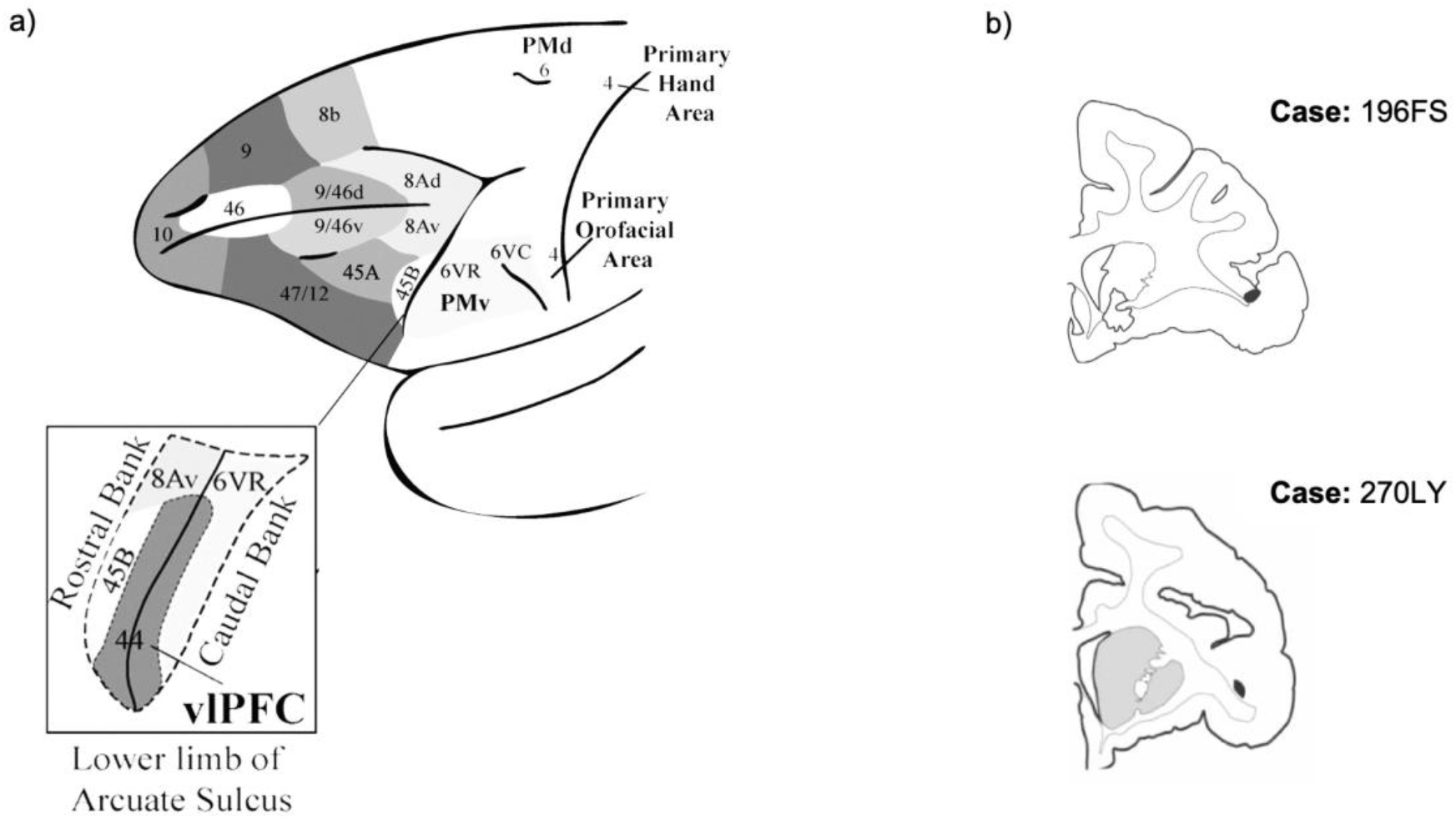
Ventrolateral Frontal Cortex. a) Schematic of ventrolateral prefrontal and ventrolateral premotor subregions (adapted from Loh et al., 2017[10]). b) Coronal sections displaying the anterograde injection sites (shaded in black) in area 44.

However, despite the posited motor-related roles of area 44, little is known about the projections of area 44 to the striatum, a crucial structure in the control of motor activity [19, 20], including speech production[21]. In the human, diffusion-weighted MRI tractography studies have provided evidence for tracts that connect area 44 to the caudate nucleus[22] and putamen[22, 23]. However, these studies do not provide information about where area 44 terminates within these structures, about the strength of the projections, or about what other corticostriatal projections they interface with. In the macaque, Choi et al (2017a)[6] and (2017b)[4] report on retrogradely labeled cells in an aggregated area 44/45 region that project to a handful of discrete areas of the striatum. Specifically, they found that more area 44/45 cells project to the dorsal than to the ventral caudate nucleus, and that few project to the ventral striatum. However, these studies did not distinguish between projection cells in area 44 and area 45 in the examined injection cases. These studies also did not examine retrogradely labeled cells from injections in the putamen – the principal motor-related region of the striatum. Furthermore, these studies did not examine anterograde injections in area 44, which are necessary to illustrate the full innervation zone throughout the striatum.

The present study sought to more fully elucidate the corticostriatal projections of area 44. First, we used anterograde tracer injections in area 44 to map its terminal field zone in the striatum. Second, we used retrograde tracer injections in the striatum to quantify the strength of area 44 projections in different parts of its innervation zone. Finally, we examined the convergence of how area 44 projections with projections from non-motor and motor related frontal regions. Of specific interest was the extent to which area 44 projection strength tracks with vlPFC and area 6VR projection strengths.

## MATERIALS AND METHODS

### Overview

First, we mapped the three-dimensional striatal innervation zone of area 44 using anterograde tracer injections. To do so, we injected an anterograde tracer into area 44 in two different macaques (**Fig. 1b)**.

Following immunohistochemistry and tissue processing, we outlined each injection site’s terminal fields in the striatum. In order to compare the terminal field locations of the different injections, we registered the terminal fields from each injection into a standardized macaque striatum [24].

Second, four retrograde tracer injections were placed into different area 44 innervation regions of the striatum to quantify the strength of the area 44 projection to each region and to evaluate area 44 convergence with other vlPFC subregions and with premotor area 6VR. Two additional retrograde injections were placed outside the area 44 innervation zone as control cases. To quantify input strength, we counted the number of retrogradely labeled cells in each frontal cortex subregion.

### vlPFC Definition

Definitions of which subregions comprise the macaque vlPFC vary. The core vlPFC regions – those most consistently included in definitions of the vlPFC across macaque anatomy research groups– are areas 12/47 (rostral and lateral) and 45[25–29]. In addition, some groups commonly include area 46v[25, 29–31], area 12/47O[26, 28], and/or area 44 in their definition of vlPFC[11, 32–35]. Definition of the human vlPFC is more consistent, and typically consists of areas 12/47, 45 and 44[36, 37]. In order to best facilitate future translation to the human, here we define the macaque vlPFC as comprising areas 12/47 (rostral, lateral and orbital), 45 (45A and 45B) and 44. Importantly, recent comparative cytoarchitectonic studies have established homologies between the macaque and human area 12/47[27], area 45[27] and area 44 subregions[11, 13].

### Surgery and Tissue Preparation

Eight adult male macaque monkeys (three Macaca mulatta, three Macaca nemestrina and two Macaca fascicularis) were used for anatomical tract-tracing experiments. All experiments were performed in accordance with the Institute of Laboratory Animal Resources Guide for the Care and Use of Laboratory Animals and approved by the University Committee on Animal Resources at University of Rochester. Details of the surgical and histological procedures have been described previously [24]. Monkeys were tranquilized by intramuscular injection of ketamine (10 mg/kg). A surgical plane of anesthesia was maintained by intravenous injection of pentobarbital (initial dose of 20 mg/ kg, i.v., and maintained as needed). Temperature, heart rate, and respiration were monitored throughout the surgery. Monkeys were placed in a David Kopf Instruments (Tujunga, CA) stereotaxic, a midline scalp incision was made, and the muscle and fascia were displaced laterally to expose the skull. A craniotomy (~2–3 cm^2^) was made over the region of interest, and small dural incisions were made only at recording or injection sites. In some animals, to guide deep cortical injections, serial electrode penetrations were made to locate the anterior commissure as described previously [38]. The absence of cellular activity signaled the area of fiber tracts, i.e., the corpus callosum, the internal capsule, and the anterior commissure. Additional recordings were performed to determine the depth of the injection sites. Accurate placement of tracer injections was achieved by careful alignment of the injection cannulas with the electrode. For more recent experiments, we obtained magnetic resonance images to guide our injection sites.

Tracers (40–50 nl, 10% in 0.1 mol phosphate buffer (PB), pH 7.4) were pressure injected over 10 min using a 0.5 μl Hamilton syringe. Tracers used for the present study were Lucifer Yellow (LY), Fluorescein (FS) conjugated to dextran amine (Invitrogen), or wheat germ agglutinin conjugated with horseradish peroxidase (WGA) (Sigma-Aldrich). After each injection, the syringe remained in situ for 20–30 min. After a survival period of 12–14 days, monkeys were again deeply anesthetized and perfused with saline, followed by a 4% paraformaldehyde/1.5% sucrose solution in 0.1 mol PB, pH 7.4. Brains were postfixed overnight and cryoprotected in increasing gradients of sucrose (10, 20, and 30%)[24]. Immunocytochemistry was performed on one in eight free-floating 50 μm sections to visualize LY, FS, or WGA tracers, as previously described [39].

### Anterograde Experiments

For each injection case, dark field light microscopy under 1.6×, 4×, and 10× objectives was used to locate and characterize area 44 terminal fields in the striatum (**Fig. 2**).

**Figure 2:**
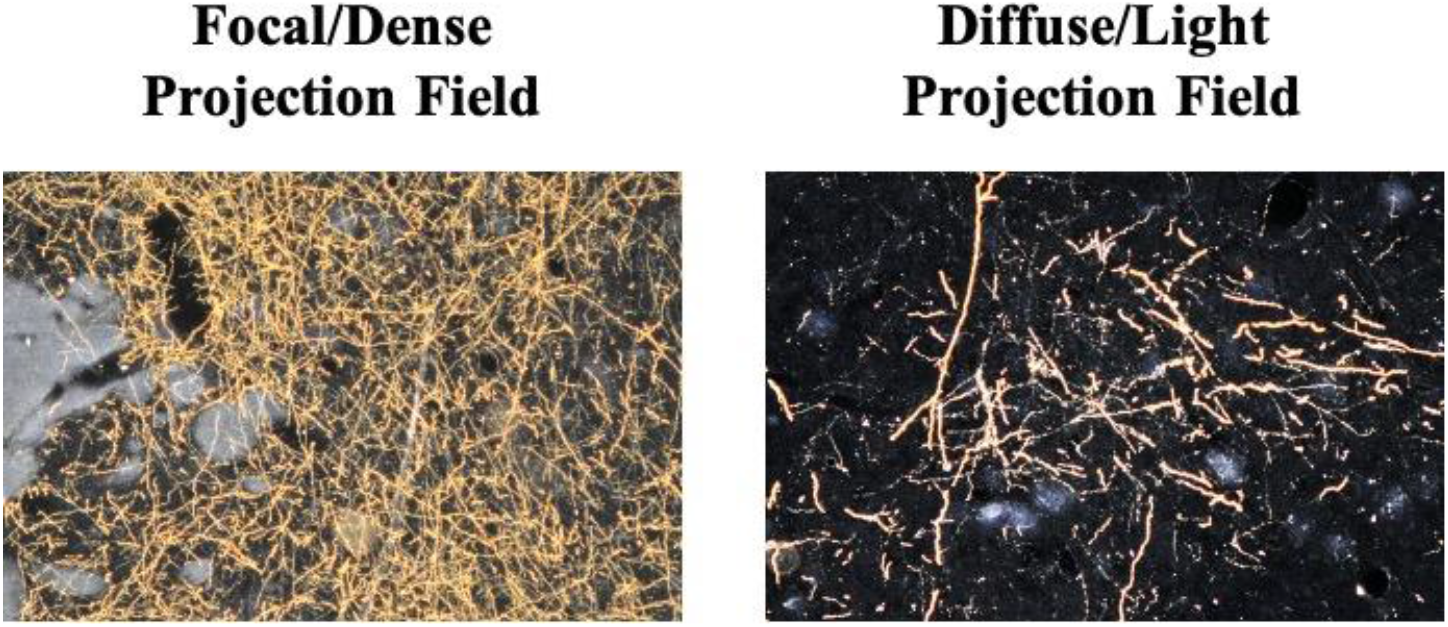
Anterogradely Labeled Terminal Fields. Photomicrographs at 10x magnification displaying densely distributed terminal field fibers and diffuse/lightly distributed terminal field fibers.

Neurolucida software (MicroBrightField) was used to chart outlines of terminals fields in the striatum in one in eight sections throughout the striatum. All thin, labeled fibers containing boutons were charted. Thick fibers without clear boutons were assumed to be passing fibers and not charted. We categorized the terminal fields into three categories: “fibers”, “lightly distributed terminal fields”, and “densely distributed terminal fields”. Individual fibers that were sparsely distributed and individually traceable were labeled as “fibers” and individually drawn. More condensed patches of fibers, but for which individual fibers within the patch could still be discerned, were labeled as “lightly distributed terminals”; the outer contours of these patches were outlined. Patches of heavily condensed groups of fibers that could not be discerned individually and which were visible at 1.6× with discernible boundaries were labeled as “densely distributed terminals”; the outer contours of these patches were outlined. Adjacent patches of dense terminals visible at 1.6x that were clearly surrounded by a less dense area, but visualized at 4x, were connected and considered as one object. Isolated patches were treated as individual objects. Boundaries for each terminal field patch were checked for accuracy at high magnification (10x).

### Transposing Cases into a Reference Model

For each case, a stack of 2-D coronal sections was created from its Neurolucida chartings. This stack was imported into IMOD (Boulder Laboratory for 3D Electron Microscopy) [40]. To merge several cases together to facilitate their comparison, we developed a reference model from one animal by sampling one in four sections (at 200-μm intervals) throughout the entire brain, using frozen and Nissl-stained sections. Data from each case were then transposed into the reference brain using landmarks of key internal structures. Following the transposition of projection terminal chartings from each case, every fiber and contour placed in the reference model was checked with the original for medial/lateral, dorsal/ventral, and anterior/posterior placement and relative size. This ensured that the chartings from each case were accurately placed with respect to their position and the proportion of the striatum they occupied.

### Retrograde Cell Experiments

Bright field light microscopy under 10X objective was used to identify retrogradely labeled frontal cortical cells. For each case, labeled cells were quantified in 37 frontal cortical subregions (using StereoInvestigator-MicroBrightField in one in twenty-four sections). To ensure complete sampling of each subregion area and to avoid double-counting, we marked each labeled cell inside of a square counting frame that was moved systematically throughout the area of each subregion on each section. A labeled cell was identified by the presence of punctate staining in the cell body. Cytoarchitectonic areal boundaries were determined using the atlas by Paxinos et al. (2000)[41].

### Analysis

To account for variability in tracer uptake and transport between different retrograde injection cases, we calculated the percentage of total labeled cells that projected from each frontal cortical subregion to each injection site (i.e. percent of total frontal cortical input). We used this metric, rather than the total number of labeled cells from each frontal cortical subregion, to compare across injection cases. This facilitated normalized comparison across injections and animals. For each injection site, we ordered the inputs from highest percent input to lowest percent input, and calculated the cumulative percent input strength at each subregion. All subregions that contributed to 90% of the cumulative frontal cortical input were considered to provide “strong” input to the striatal injection site. Subregions beyond the 90% cutoff point contributed negligible input (1% or less of the total frontal cortical input) to the injection site.

## RESULTS

### Anterograde Tracing

Collectively, area 44 projections terminate in the rostral central and ventrolateral parts of the striatum (**Figs. 3-4**).

**Figure 3:**
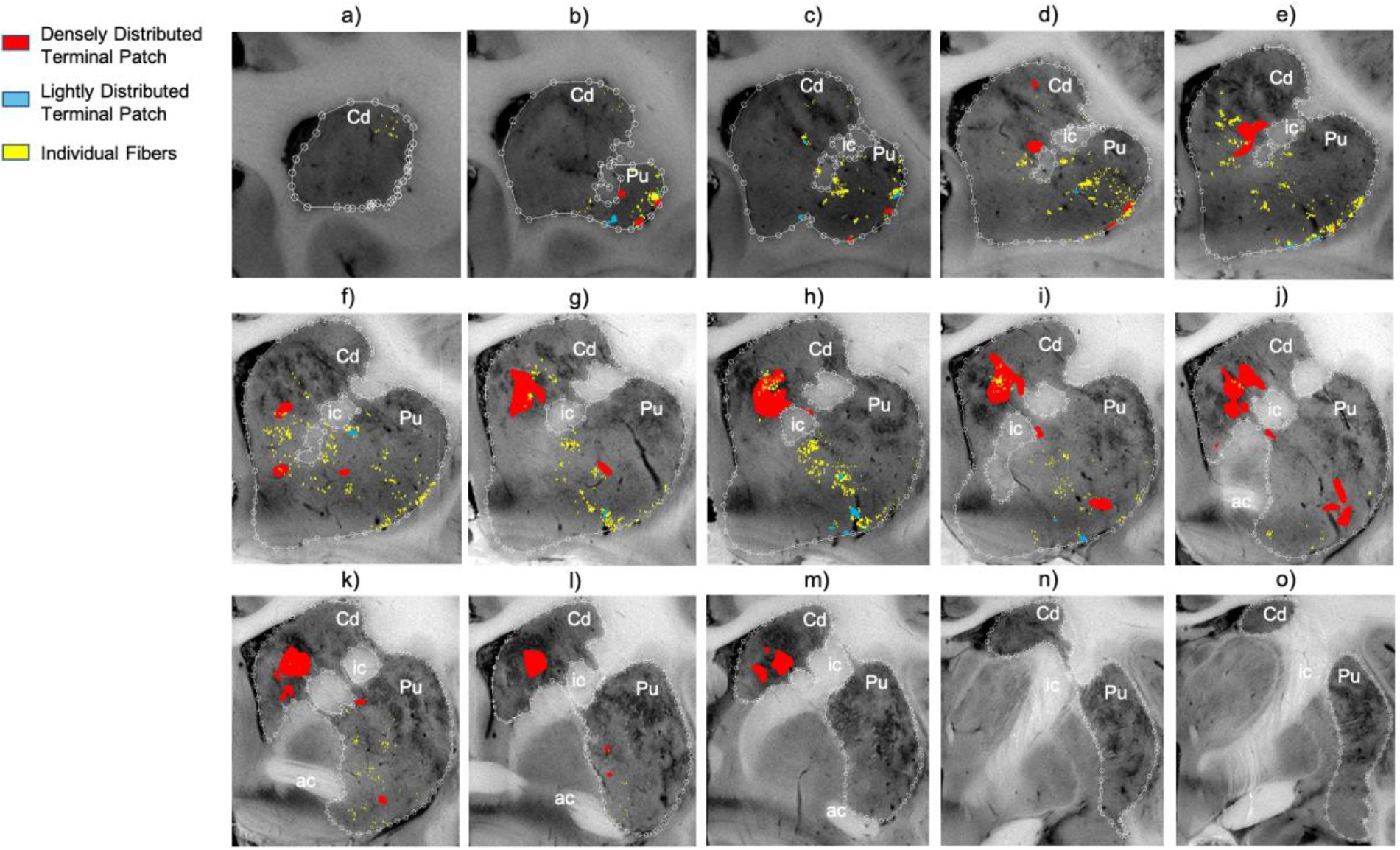
Distribution of Focal and Diffuse Area 44 Projection Fields. a-o) 2D coronal slices through the striatum depicting the densely distributed terminal patches (focal fields), and lightly distributed terminal patches and individual fibers (diffuse fields) of the area 44 injection sites. Slices proceed from rostral (left) to caudal (right). ac, Anterior commissure; Cd, caudate nucleus; ic, internal capsule; Pu, putamen nucleus.

**Figure 4:**
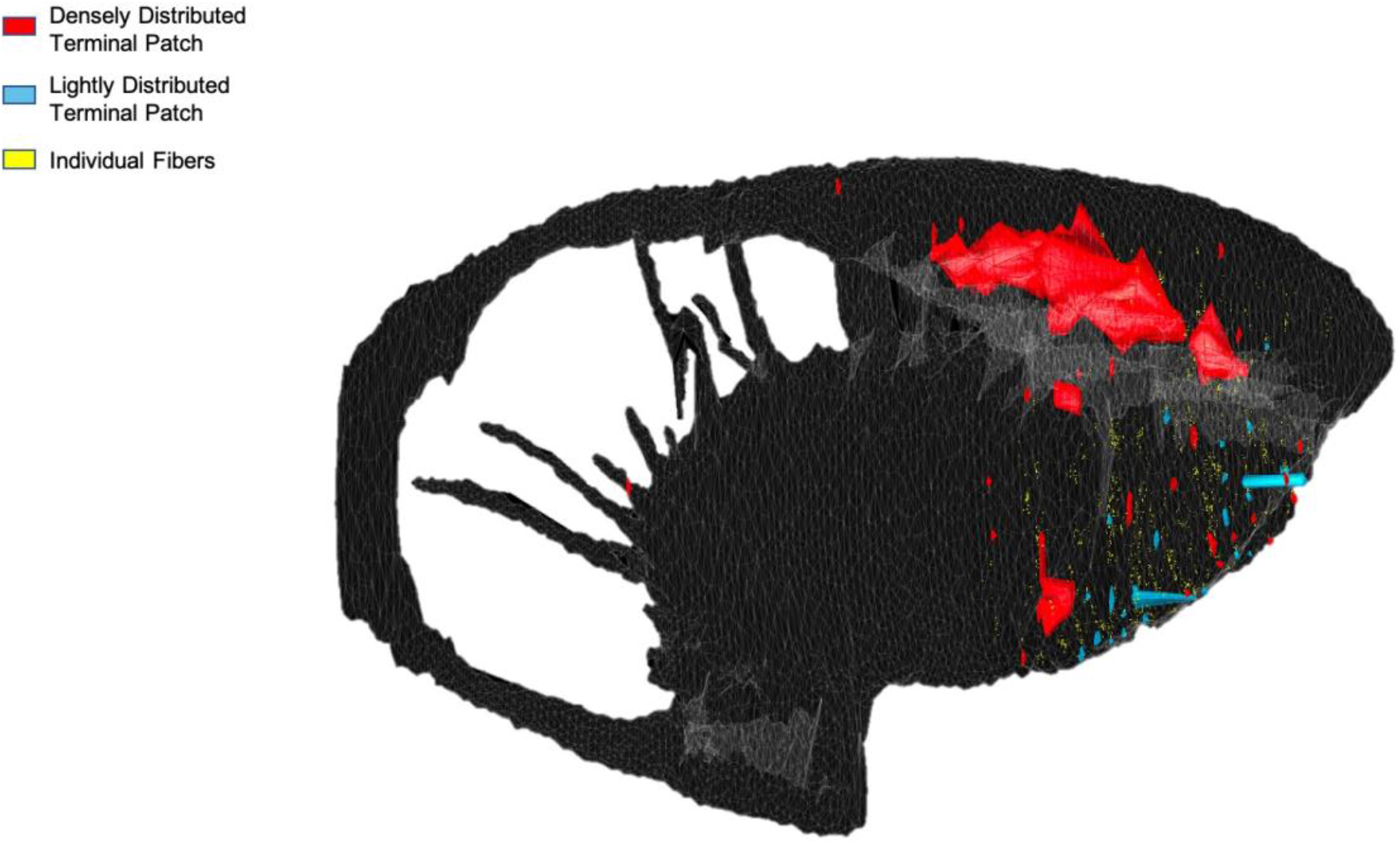
Area 44 Projections in 3D Space. Three-dimensional rendering (sagittal view) of the area 44 terminals in the striatum.

The terminal field forms a diagonal band from the central caudate nucleus to the ventrolateral putamen. Along the dorsal-ventral axis, the area 44 terminal band is dorsal to – and thus does not include – the ventral striatum. Furthermore, except at very rostral levels (**Fig. 3a-d**), it is ventral to – and does not include - the dorsolateral caudate nucleus or dorsal putamen. Along the rostral-caudal axis, area 44 terminals are concentrated rostral to the caudal end of the anterior commissure; there are few terminals caudal to this point. There are also few terminals in the rostral striatal pole. Consistent with previous studies [24], there are two area 44 terminal labeling patterns in the striatum: a focal, or dense, projection field and a diffuse, or light, projection field (**Fig. 3**). Dense terminal fields from area 44 are primarily located in the central caudate nucleus (**Fig. 3d-m**), with some patches in the rostral ventral putamen (**Fig. 3b-d**) and in the posterior ventral putamen (**Fig. 3i-l**). Lightly distributed terminal patches and individual fibers are also present in the rostral dorsal caudate nucleus (**Fig. 3a-d**) and rostral central putamen (**Fig. 3b-f**).

We compared the location of terminal fields between the more rostral and more caudal injections (196FS and 270LY respectively) (**Fig. 5**). Both injection sites result in innervation of a diagonal band of similar orientation and of similar dorsal-ventral/medial-lateral/rostral-caudal placement (both include the central caudate nucleus and the ventral putamen). However, there is a rostro-caudal topography: the rostral injection projects more substantially to the caudate nucleus, and the caudal injection projects more substantially to the putamen. Note: while the position of terminal fields is clear for both cases, tracer uptake and transport is stronger for the rostral injection site, resulting in more dense terminal patches than for the caudal injection site.

**Figure 5:**
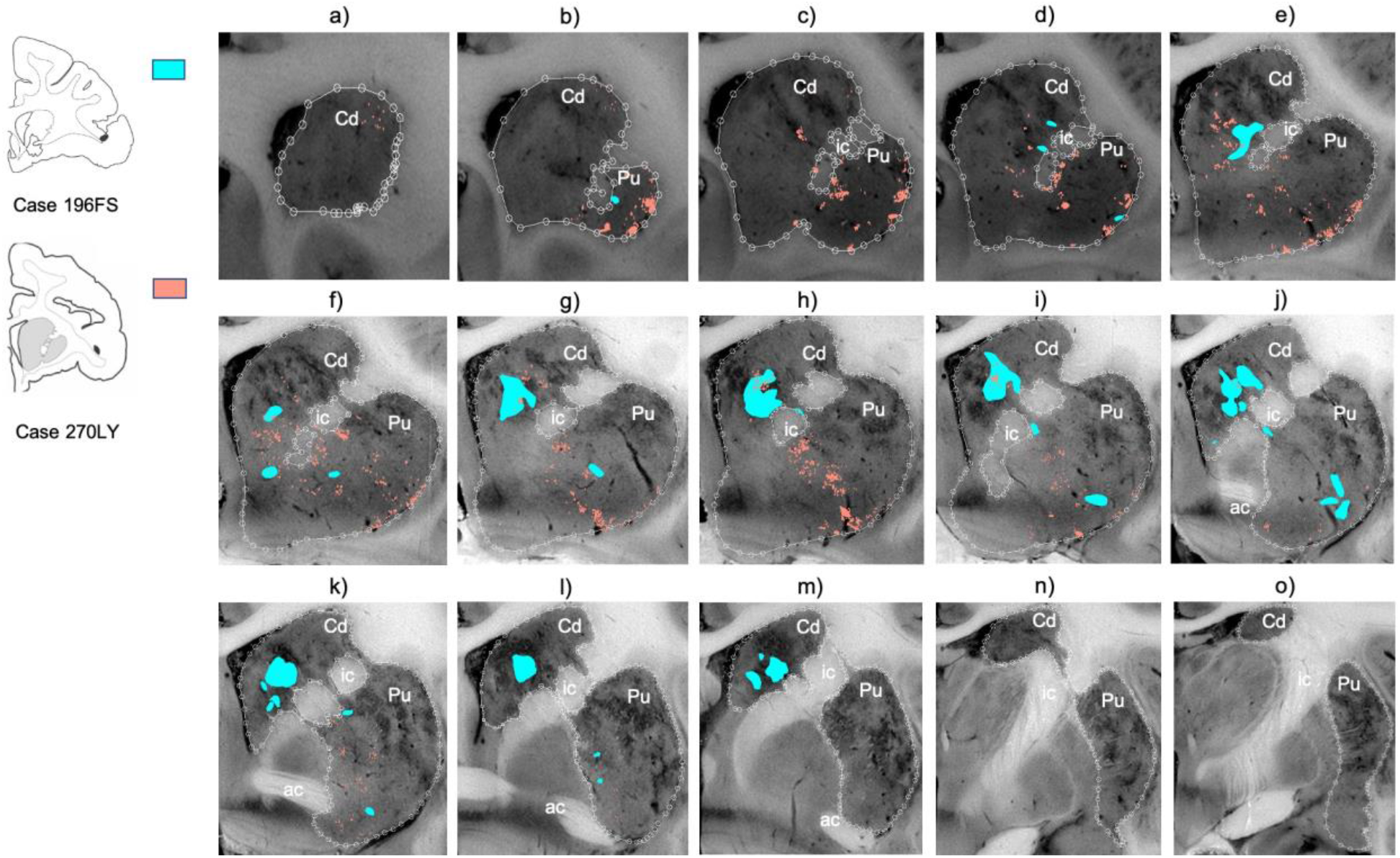
Comparing Terminal Fields from Different Injections in Area 44. a-o) 2D coronal slices through the striatum, comparing the terminal fields from the two area 44 injection sites. Slices proceed from rostral (left) to caudal (right). Densely and lightly distributed terminal fields are depicted in the same color. ac, Anterior commissure; Cd, caudate nucleus; ic, internal capsule; Pu, putamen nucleus.

### Retrograde Tracing

#### Area 44 Projection Strengths

We quantified the percent of total frontal cortical input contributed by area 44 at four sites within the area 44 innervation zone (cases 170FS, 252WGA, 253WGA, and 39WGA) and at two sites outside the area 44 innervation zone (cases 35LY and 38LY) (**Fig. 6**). As expected from the anterograde results, the fewest retrogradely labeled cortical cells were seen following injections in the striatal areas that contained the fewest anterogradely labeled terminal fields– the rostral pole and ventral striatum - (0.2% and 1.2% of total frontal cortical input, respectively). The strongest input from area 44 compared to other cortical inputs resulted from the caudal ventral putamen injection site (10.8% of total frontal cortical input). In contrast, the dorsal caudate nucleus, ventral caudate nucleus, and ventrolateral putamen injection sites resulted in lower, but similar input strength (4.5%, 5.0%, and 4.7% of total frontal cortical input, respectively).

**Figure 6:**
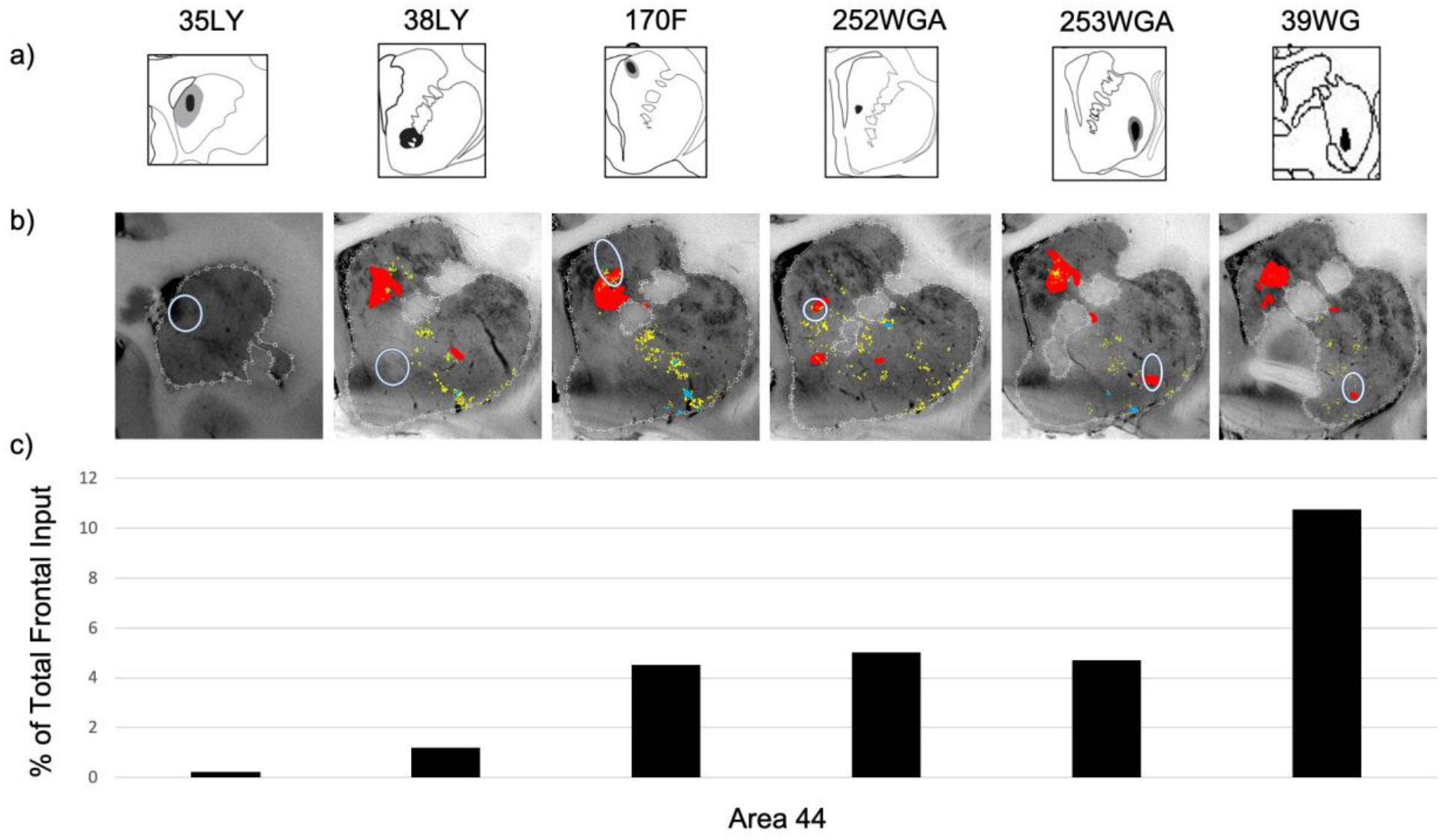
Quantifying Area 44 Input Strength. a) Striatal retrograde injection sites (coronal view); b) The injection sites (white circles) overlaid on the anterogradely labeled area 44 terminal fields; c) For each striatal injection site, the percentage of all frontal cortical retrogradely labeled cells that were in area 44.

The anterograde and retrograde data displayed robust correspondence (**Fig. 6**). Retrograde injections whose location did not overlap with anterogradely labeled area 44 terminals displayed the weakest area 44 input. In contrast, retrograde injections whose location overlapped with dense anterogradely terminal fields from area 44 terminals also displayed the most area 44 retrogradely labeled cells.

#### Convergence of Projections from Area 44 with those from Area 6VR and the vlPFC

At injection sites in the putamen component of the area 44 innervation zone, strong area 44 input was accompanied by strong area 6VR input, but negligible vlPFC or other prefrontal input (**Fig. 7**). Out of the 37 frontal cortical areas, area 44 and area 6VR were the second and first strongest inputs, respectively, at the caudal ventral putamen injection site, and were the fifth and fourth strongest inputs, respectively, at the ventrolateral putamen site. The majority of the remaining input at these sites was contributed by motor-related regions, including the motor cingulate area (area 24c), the pre-supplementary and supplementary motor areas (area 6M), and ventral premotor areas 6VC and ProM.

**Figure 7:**
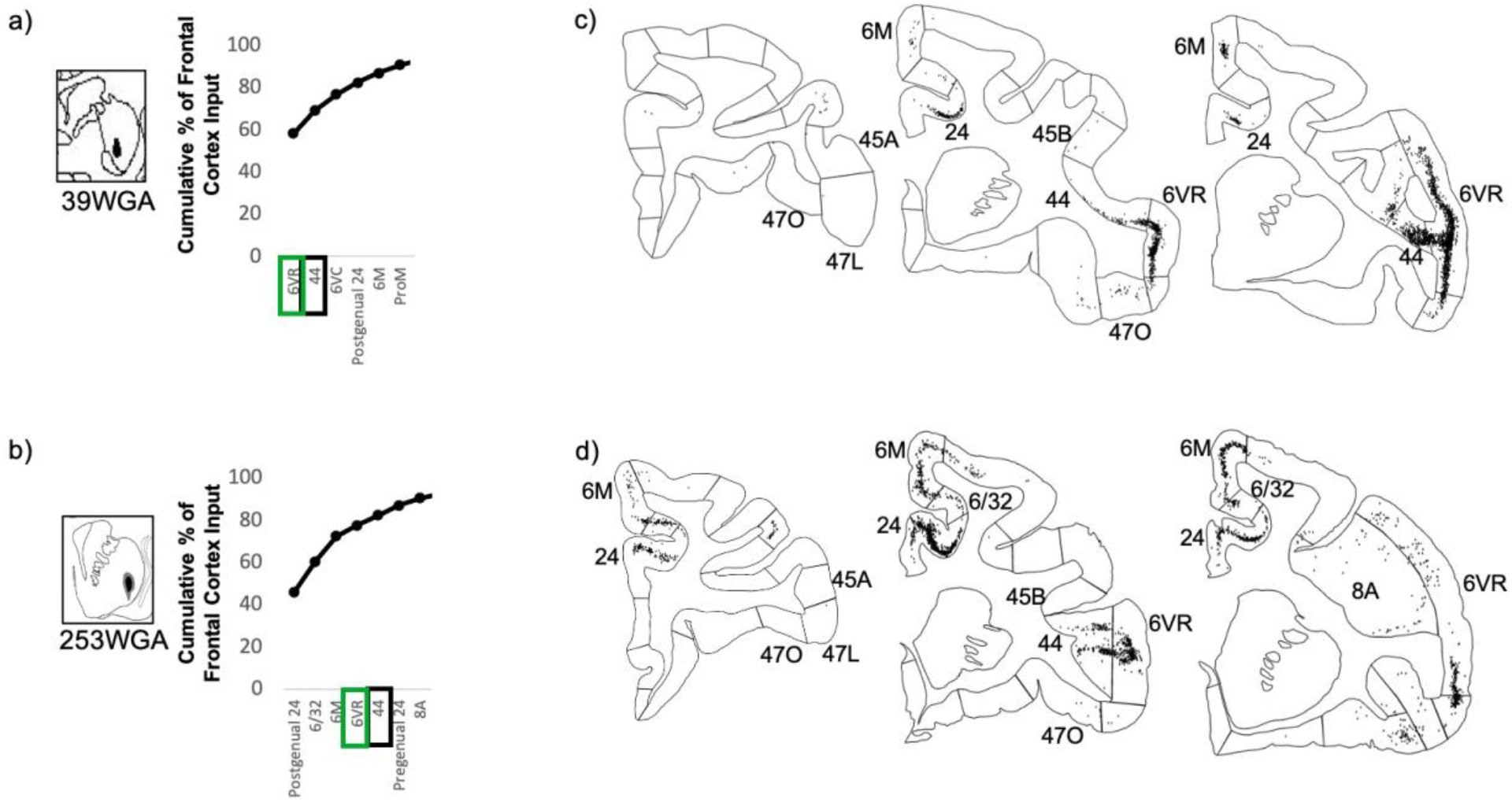
Striatal injection sites with strong area 44 input and strong area 6VR input. a-b) The cumulative percent of total frontal cortex input contributed by each frontal cortical region to the injection site, up to 90% of cumulative input. Area 6VR outlined in green. Area 44 outlined in black. c-d) For each striatal injection site, representative coronal slices displaying retrogradely labeled cells.

Conversely, at injection sites in the caudate nucleus component of the area 44 innervation zone, strong area 44 input was accompanied by strong vlPFC and other prefrontal cortical input, but negligible 6VR input (**Fig. 8**). In both the ventral and dorsal caudate nucleus, strong area 44 input was accompanied by strong input from vlPFC area 47R, dorsolateral prefrontal areas 46V and 9L, dorsomedial prefrontal area 9/32, and cingulate area 24. In the ventral caudate nucleus specifically, there was also strong input from vlPFC areas 47L, 470 and 45A, orbitofrontal areas 11, 13 and 14O and ventromedial prefrontal area 25. And in the dorsal caudate nucleus specifically, there was also strong input from vlPFC area 45B, frontal pole areas 10L and 10M, frontal eye field areas 8B and 8A, and premotor area 6DR. Out of the 37 frontal cortical subregions, area 44 was the sixth strongest input at the ventral caudate nucleus site, and was the tenth strongest input at the dorsal caudate nucleus site.

**Figure 8:**
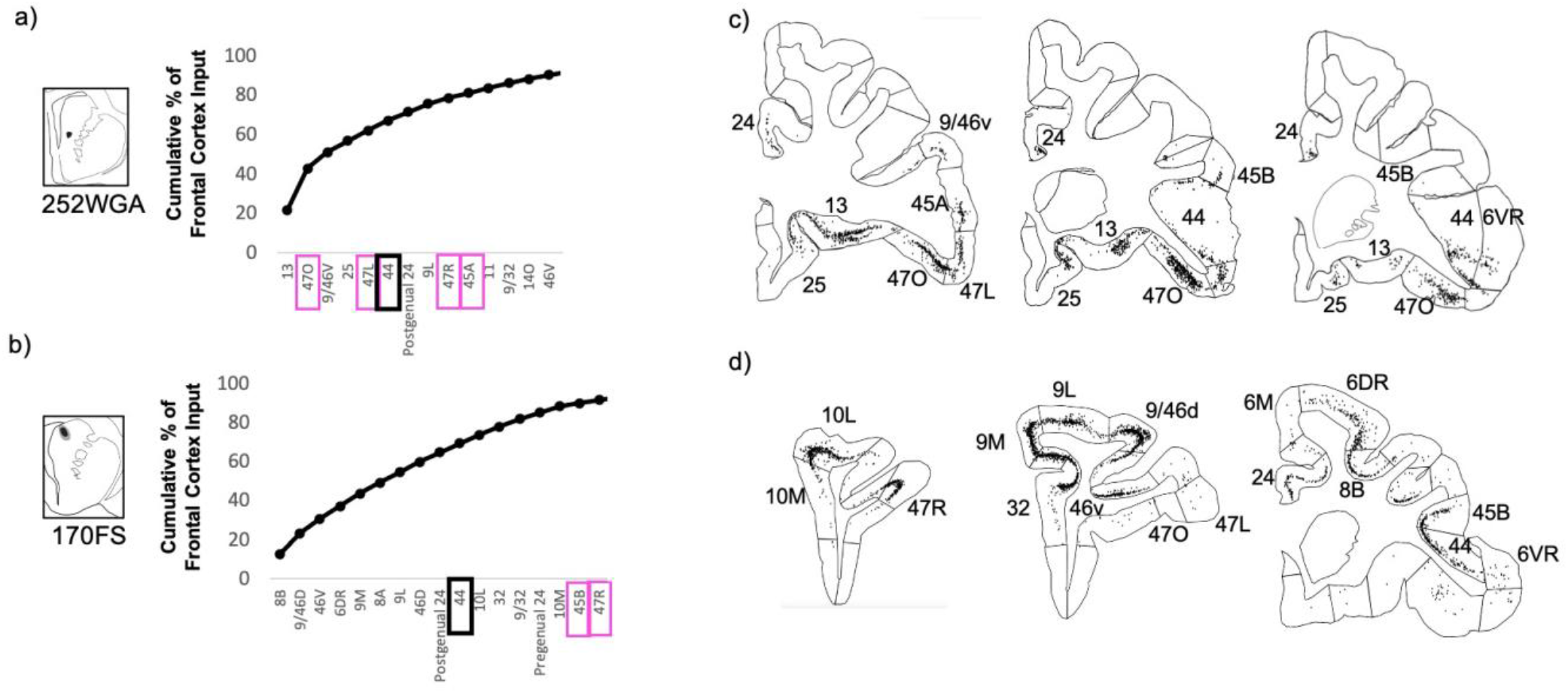
Striatal injection sites with strong area 44 input and strong vlPFC input. a-b) The cumulative percent of total frontal cortex input contributed by each frontal cortical region to the injection site, up to 90% of cumulative input. Area 44 outlined in black. Other vlPFC subregions outlined in pink. c-d) For each striatal injection site, representative coronal slices displaying retrogradely labeled cells.

At the injection sites outside of the area 44 innervation zone, negligible area 6VR input accompanied the negligible area 44 input (**Fig. 9**).

**Figure 9:**
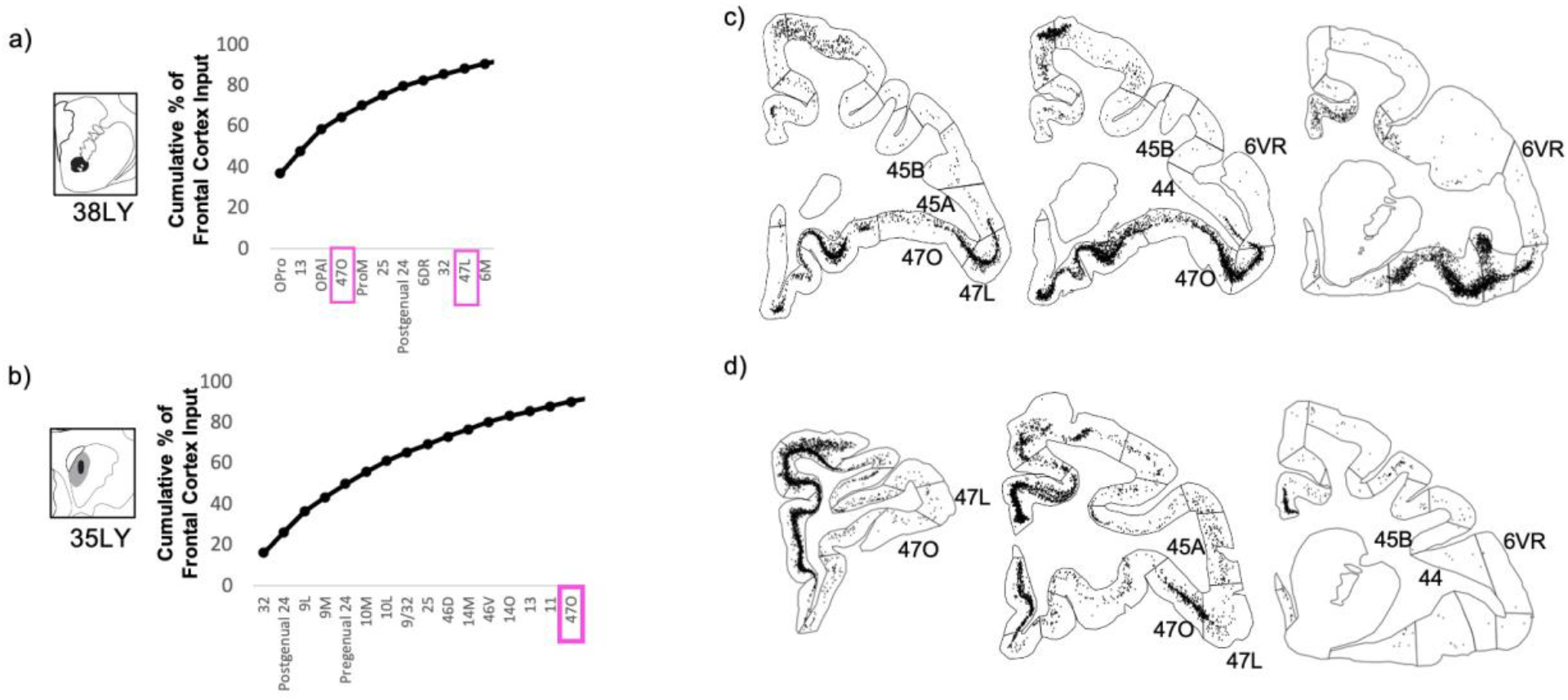
Striatal injection sites with negligible area 44 input. a-b) The cumulative percent of total frontal cortex input contributed by each frontal cortical region to the injection site, up to 90% of cumulative input. vlPFC subregions outlined in pink. c-d) For each striatal injection site, representative coronal slices displaying retrogradely labeled cells.

### Limitations

Tract-tracing studies have several inherent limitations. First, there is the potential for tracers to be picked up by fibers of passage, especially in striatal fascicles, and thus label cells whose projections do not terminate at the injection site. Furthermore, since WGA can be transported transynaptically to some extent after 48-72 hours, some labeled cells in WGA cases in this study may represent polysynaptic connections. However, several reports indicate that WGA transsynaptic staining may primarily be limited to cells in close proximity to monosynaptically labeled cells[34, 42, 43]. This could result in inflated cell counts within WGA cases in the present study. However, given the small probability of long distance polysynaptic transport coupled with our metric of interest being percent – rather than absolute number – of labeled cells, this should have little to no effect on cross-region comparisons. Another limitation is that the alignment of the retrograde injection sites with the anterogradely defined area 44 innervation zone is imperfect. This is particularly evident for injection 170FS, which contains some of the area 44 innervation zone, but a larger striatal area that is outside the innervation zone. As such, injection sites cannot be said to be completely inclusive or exclusive of the area 44 innervation zone. Finally, it is important to note that the retrograde injection sites analyzed here do not represent a full sampling of the area 44 innervation zone, and that injections at other sites not sampled here may reveal different profiles of input strengths.

## DISCUSSION

### Summary

Ventrolateral frontal area 44 has been implicated in motor functions in both humans and macaques, including speech and vocalizations[7–12]. However, the projections from area 44 to the striatum – a crucial structure in motor circuits[19, 20] – have received little study. In particular, given the posited role of area 44 in facilitating prefrontal control over orofacial motor activity[10], it was of interest to investigate how corticostriatal projections from area 44 may converge with those from the adjacent ventrolateral prefrontal cortex and ventrolateral orofacial premotor area 6VR. Here, we used anterograde and retrograde tract-tracing in macaques to elucidate the three-dimensional innervation zone of area 44 projections to the striatum, quantify the strength of these projections, and evaluate their convergence with corticostriatal projections from vlPFC and area 6VR. First, we found that area 44 projections terminate in the rostral central and ventrolateral parts of the striatum, in a diagonal band dorsal to the ventral striatum that spans from the central caudate nucleus to the ventrolateral putamen. Terminal fields from a rostral area 44 injection were found primarily in the caudate nucleus, whereas terminal fields from a caudal area 44 injection site were found primarily in the putamen. Second, we found that input from area 44 accounted for between 4.5% and 10.8% of total frontal cortical at sites sampled within its innervation zone. While the largest focal projections from area 44 were found in the caudate nucleus, area 44 input accounted for its greatest relative proportion of total frontal cortical input in the ventral putamen at the level of the anterior commissure, due to the small amount of input from almost all other frontal cortical regions to this site. Third, we found that in the putamen component of the area 44 innervation zone, strong area 44 input was accompanied by strong area 6VR and other motor-related input from the motor cingulate area, pre-SMA, and SMA. Conversely, in the caudate nucleus component of the area 44 innervation zone, strong area 44 input was accompanied by strong vlPFC input, as well as other non-motor-related prefrontal input. Overall, the anatomical positioning of area 44 corticostriatal projections support its role in motor circuitry, and also suggests a role in non-motor circuits involving the caudate nucleus and prefrontal cortex. This is consistent with prior literature that has conceptualized area 44 as an interface between motor and non-motor functional domains. The present results suggest that this may be facilitated by a more rostral subdivision of area 44 that primarily projects to the caudate nucleus and converges with non-motor prefrontal projections, and a more caudal subdivision that projects primarily to the putamen and converges with motor-related projections.

### Area 44 Striatal Innervation Zone

The diagonal band that characterizes the area 44 innervation zone in coronal sections is consistent with the structure of striatal innervation zones from other cortical regions[1]. Furthermore, its dorsal-ventral/medial-lateral location aligns with the general topographic organization of the striatum [44–46]. More ventral and medial frontal cortical areas innervate bands that span more of the ventromedial striatum, whereas more dorsal and lateral frontal cortical areas innervate bands that span more of the dorsolateral striatum. Ventrolateral prefrontal areas, located between these ventromedial and dorsolateral extremes, have been shown to innervate diagonal bands that span more of the central striatum[3, 47–50]. The location of the area 44 diagonal band resembles those from these other ventrolateral frontal areas. In addition, consistent with previous work [24], we observed two area 44 terminal labeling patterns in the striatum: a focal, or dense projection field, and a diffuse, or light projection field. Dense terminal fields from area 44 are primarily in the central caudate nucleus, with some in the rostral ventral putamen and more posterior ventral putamen as well. Lightly distributed terminal patches and individual fibers are also present in the rostral dorsal caudate nucleus and rostral central putamen. Overall, this detailed characterization of the area 44 innervation zone *within* the striatum builds upon prior diffusion-weighted MRI tractography studies in humans that demonstrated tracts connecting area 44 *to* the striatum[22, 23].

### Quantitative Strength of Area 44 Corticostriatal Projections

These results expand upon the limited prior data on the strength of corticostriatal projections from area 44. Our findings of negligible area 44 innervation of the ventral striatum are consistent with the findings from Choi et al. (2017b)[4] that aggregated area 44/45 projects minimally to this region. In the caudate nucleus, Choi et al. (2017a)[6] reported that the density of aggregated area 44/45 cells that project to the dorsomedial caudate nucleus is higher than the density of such cells that project to the ventral caudate nucleus. However, some of this difference may have been attributable to differences in tracer uptake and transport between the dorsal and ventral caudate nucleus injection cases. Here, we find that area 44 contributes a similar proportion of the total frontal cortical input at both sites (4.5% of frontal cortical input in the dorsomedial caudate nucleus, and 5.0% of frontal cortical input in the ventral caudate nucleus). As such, regardless of the absolute number of area 44 cells that project to each site, area 44 likely has a similar degree of computational influence, relative to other frontal cortical regions, at both sites.

We also demonstrate the input strength of area 44 in the putamen component of its innervation zone. At the ventrolateral putamen injection site just rostral to the anterior commissure, area 44 projections comprise 4.7% of the total frontal cortical input, similar to the level observed in the dorsomedial and ventral caudate nucleus. However, at the ventral putamen site at the caudal end of the anterior commissure, area 44 input constitutes 10.8% of the total frontal cortical input. These data suggest that the relative influence of area 44 on striatal processing is not uniform in all parts of its innervation zone, and that area 44 may have its greatest relative influence on striatal processing in the ventral putamen at the caudal end of the anterior commissure.

### Convergence of Area 44 Corticostriatal Projections with those from vlPFC and area 6VR

Overall, the corticostriatal projections from area 44 shared features of those from both the vlPFC and orofacial premotor area 6VR. Area 44 input strength tracked more closely with input strength from area 6VR in the putamen, where both projected strongly and where vlPFC projections were negligible. Conversely, area 44 input strength tracked more closely with input strength from vlPFC in the caudate nucleus, where both projected strongly and where area 6VR projections were negligible. In this way, the area 44 corticostriatal projection is distinct from the corticostriatal projection of both the adjacent prefrontal and premotor cortices - possibly mediated by rostral and caudal subregions with differing projection profiles. The mixed prefrontal/premotor properties of the area 44 corticostriatal projection mirror that of its corticocortical connections: while area 44 is more strongly connected to premotor cortical areas than the adjacent vlPFC is[17, 51], it is also more connected to prefrontal areas than the adjacent premotor cortex is[10]. As such, area 44 is anatomically positioned to interface with the executive and motor components of orofacial motor control via both its corticocortical and corticostriatal connections,. The strong area 44 corticostriatal input to the ventral putamen injection sites was also accompanied by strong input from dorsomedial motor-related regions, including area 6M (the presupplementary and supplementary motor areas). In the human, area 44, area 6M, and putamen have been implicated as central structures in a motor inhibition network[52–56]. Our observation here that the ventral putamen receives strong input from both area 44 and area 6M is suggestive of a possible anatomical underpinning for the interaction of these structures during motor inhibition.

Finally, we observed that area 44 corticostriatal projections converged with those from frontal cortical areas that area 44 is not known to have strong corticocortical connections with[17]. This was particularly evident in the caudate nucleus, where strong area 44 projections converged with strong projections from orbitofrontal cortex, ventromedial prefrontal cortex, and dorsolateral prefrontal cortex. These interfaces may suggest an as of yet underappreciated role of area 44 in non-motor functions.

## ACKNOWLEDGMENTS

This work is funded by NIMH grant 2R01MH045573-29A1 to SNH and NIDA grant 1F32DA048580-01A1 to CK. The authors report no conflicts of interest.

